# Plant species and floral traits shape arthropod communities in restored prairies more than neonicotinoid contamination

**DOI:** 10.1101/2025.04.23.650261

**Authors:** Jonathan Tetlie, Alexandra Harmon-Threatt

## Abstract

Agricultural practices are significant drivers of biodiversity loss, leading to reductions in ecological function and services across regions. To mitigate these effects, habitat restorations within agroecosystems have gained prominence as a strategy to enhance ecological stability and increase biodiversity. However, the pervasive use of neonicotinoids in agroecosystems has raised concerns about their impact on restored habitats in agriculturally dominant regions. This study investigates the multi-trophic effects of neonicotinoid contamination on arthropod community structure and plant fitness in early prairie restoration ecosystems. Using a manipulated field experiment and structural equation modeling, we assessed how neonicotinoid exposure influences arthropod abundances across different feeding guilds, abundance interactions between feeding guilds, and plant fitness, measured by seed set and aboveground biomass. Our findings reveal that neonicotinoid contamination does not uniformly affect arthropod feeding guilds; instead, floral characteristics predominantly drive community composition and interactions.

While neonicotinoid exposure significantly increased pollinator visitation and decreased omnivore abundance in one of our models, plant species and inflorescence abundance were significantly more impactful on arthropod feeding guilds and community structure. These results underscore the importance of plant community composition in promoting prairie restoration stability and suggest that the benefits of increasing restored prairie habitats outweigh the potential risks of more critical for ecosystem resilience than mitigating neonicotinoid contamination.

## Introduction

Agricultural practices are one of the largest contributors to biodiversity loss and can drastically reduce the number of ecological niches available within a region (Dudley and Alexander, 2017; Wagner et al., 2021). To counteract this decline, the restoration of degraded habitats, particularly within agroecosystems, has emerged as a prevalent strategy for enhancing ecological stability through increased biodiversity and the development of resilient ecosystems (Tilman et al., 1998; Biondini, 2007). However, restored lands often carry a legacy of pesticide use and are commonly found near conventional agriculture, leading to frequent contamination of these habitats by agricultural inputs (Aseperi et al., 2020). This contamination could significantly impact the ability of restored areas to help improve ecosystem stability (Ruiz-Jaen and Aide, 2005; Grman et al., 2010).

Given the great number of ecological niches provided by arthropods (Longcore, 2003), the pesticide contamination of restored areas could threaten the myriad of critical ecosystem services arthropods provide by differentially impacting groups of arthropods and altering their ecological interactions. The most widely used class of insecticides globally are neonicotinoid insecticides (Humann-Guilleminot et al., 2019; Hall et al., 2022). Neonicotinoids are regularly found in restored habitats (Main et al., 2014; Hall et al., 2022), can exhibit longer environmental persistence than other conventional insecticides (Rexrode et al., 2003; Khan, 2016), and have been identified as a prominent driving factor behind insect declines (Goulson, 2013; Pisa et al., 2021; Wagner et al., 2021). Despite this, we lack a comprehensive understanding of how arthropod communities, in conjunction with floral communities within restorations, respond to neonicotinoid contamination from a systematic perspective.

The vast majority of arthropod-neonicotinoid studies are limited in their ability to address how the ubiquitous neonicotinoid contamination of restorations in agroecosystems could lead to variation in ecological community structure and subsequent ecosystem services. By focusing primarily on laboratory-based studies, using individual species or narrow taxonomic groups in isolation, much of the current work on neonicotinoid effects lacks ecologically relevant field context (Main et al., 2018; Disque et al., 2019). Isolated assays and studies on single species have identified important species-specific responses to neonicotinoids (Nauen and Elbert, 1997; Morales-Rodriguez and Peck, 2009; Kessler et al., 2015), however these isolated results fail to characterize the impacts of neonicotinoids on broader ecosystem-level impacts such as feeding guild abundances (i.e., herbivores, predators, pollinators, etc.) and multi-trophic interactions.

Differential responses to neonicotinoids within the same system could lead to alterations in community structure and subsequent changes in ecosystem services. For example, Harmon et al., (2023) observed reductions in early-season herbivore biomass and prolonged reductions in predator biomass due to community-level neonicotinoid contamination. This suggests neonicotinoid contamination has the potential to destabilize arthropod communities by affecting certain trophic levels more than others, which could subsequently change the ecosystem and agricultural services that restorations provide. Furthermore, given the wide range of possible changes in arthropod behavior, biomass, and diversity to neonicotinoid exposure, there is likely an impact on the fitness and growth of plants, which is poorly studied and similarly could have an outsized impact on the stability of restoration ecosystems (Harvey et al., 2008; Burghardt and Tallamy, 2013; Schuldt et al., 2014).

To gain a more comprehensive and system focused understanding of how neonicotinoid contamination affects arthropod communities and the ecosystem services they provide, we used a manipulated field experiment to examine questions of how ecosystem stability, measured as seed set and plant aboveground biomass, may be impacted by neonicotinoid contamination. This design allowed for greater control of confounding variables while utilizing an established natural community, therefore maintaining valuable biological relevance. Here, we ask (i) are arthropod abundances from different feeding guilds variably affected by neonicotinoid contamination, (ii) does neonicotinoid contamination alter interactions between feeding guilds, and (iii) do either of these factors impact plant fitness – characterized in this study by plant seed set and aboveground biomass. We used structural equation modeling (SEM) to disentangle the interconnected relationships between plants, arthropod feeding guild abundances, and neonicotinoids. This approach allowed us to utilize a hypothesized structure of interspecific relationships to examine the direct and indirect effects of neonicotinoids throughout the system and assess the relative importance of each connection within the system. We constructed two SEMs to look at plant aboveground biomass and seed set separately to better assess the short- and long-term establishment of prairie plants in these ecosystems (Biondini, 2007). To further examine trends and confirm findings on a different scale, we used non-parametric analysis of variance (PERMANOVA) to compare the influence of neonicotinoid contamination and plant species on arthropod family community structure. This method assessed taxonomic differences in arthropod community structure rather than feeding guild abundances. This experimental design, paired with these analytical techniques, provides a more comprehensive representation of the effects that neonicotinoid contamination have on the stability of plant-insect interactions in early prairie restoration communities.

## Methods

### Experimental Design

To test the effects of neonicotinoids on insect plant visitation, we created experimental field pots using four prairie plants commonly included in tallgrass prairie restorations (Kurtz, 2013) and are considered attractive to pollinators (Angelella and O’Rourke, 2017): *Chamaecrista fasciculata* (Fabaceae)*, Coreopsis tinctoria* (Asteraceae)*, Rudbeckia hirta* (Asteraceae), and *Monarda fistulosa* (Lamiaceae). Experimental pots consisted of 15-gallon cloth planting pots (247 Garden, Montebello, CA) that were filled with a sanitized general-purpose soil mixture (1:2:2 – Soil:Peat:Perlite) prepared by the University of Illinois College of Agricultural, Consumer, and Environmental Sciences (ACES) Plant Care Facility (Urbana, IL), and planted with the four species. Year-old seedlings for our perennial plant species that flower during their second year (*Monarda fistulosa* and *Rudbeckia hirta*) were purchased from Prairie Nursery (Westfield, WI), while annual plants were grown from seeds purchased from Prairie Moon Nursery (Winona, MN). Two treatments, clothianidin-treated and control, were replicated 20 times each at Phillips Tract natural area (Urbana, IL). Clothianidin-contaminated pots were treated with a high level of clothianidin calculated from maize application rates (80g a.i. per hectare) in the form of Arena 0.25G (Nufarm Americas Inc., Alsip, IL) (Badua et al., 2021). We used a granular insecticide as a proxy for agricultural seed coatings, which is the most prominent application technique of neonicotinoids (Hladik et al., 2018).

This experimental design captured variations in plant life history traits (annual and perennial) and bloom time. It also allowed for greater control over neonicotinoid concentration, soil composition, and soil microbiome heterogeneity to decrease confounding variables in our data. By controlling these factors and the spatial extent, we can observe treatment effects that might be less apparent at a landscape level where contamination is widespread and heterogeneous.

### Insect visitation

Insect incidence was recorded every other week starting at the beginning of flowering in late June and continued through the first week of September 2022. Each sampling period consisted of a one-minute observer acclimation period (which was previously determined to be a sufficient amount of time for insect visitation to resume), followed by five-minute observations of plant pots where the number of active inflorescences per plant, all arthropods that were physically on each plant, and the plants on which the arthropods were found were recorded.

Arthropods were identified to the lowest taxonomic group possible, and photos were taken to assist with identification (Borror et al., 1998; Thomas et al., 2002; Whitfield et al., 2013). To limit plant damage, insect voucher specimens were not collected.

### Plant fitness sampling

Aboveground biomass and average seed set were collected in late September 2022 after plants senesced. Plants from several pots were damaged by deer during the summer and were excluded from all analyses. All remaining plants from all pots were sampled for aboveground biomass.

Samples were dried at 60°C and weighed. Flower heads were collected from a random subset of pots (Appendix S1: Table S1). Only flowers from plants that were not deer damaged and had fully senesced were collected. Three flower heads were taken from each plant, and the average per plant was used for analysis. Flower heads were dried, individually threshed, and the seeds from each head were manually counted. *Monarda fistulosa* did not flower throughout the experiment, therefore seed set was not assessed for this species.

### Statistical Analysis - SEM

To analyze community-level responses between arthropod feeding guild abundances, plant fitness outcomes, and clothianidin soil contamination, we employed structural equation modeling (SEM). Given the difference in sampling groups between aboveground biomass and seed set as mentioned earlier, two SEMs were constructed. Furthermore, these plant fitness metrics could be differentially affected by feeding guilds – which utilize different portions of the plant (Nemec et al., 2014) – and could ultimately affect the long-term establishment and proliferation of prairie restorations. Separate datasets were created for each SEM so that only complete samples for aboveground biomass and seed set were included for each SEM analysis.

Traditional SEMs – using *R* packages such as *lavaan –* rely exclusively on variance-covariance matrices between variables, do not account for non-normality of error variances between variables, cannot account for clustering of samples (random effects), and require many degrees of freedom for complex models. Because our variables of interest were sampled from multiple plant species and contain count data, which require different error distributions, we employed the *piecewiseSEM R* package (Lefcheck, 2016). Piecewise SEM combines separate generalized linear mixed-effects models into a single causal network and can accommodate both random-effects structures and non-normal distributions (Shipley, 2009).

We constructed a hypothesized structural equation model based on assumed relationships between variables (Appendix S1: Figure S1). From this, generalized linear mixed-effects models (GLMMs) were constructed based on these hypothesized relationships and later used in the piecewise SEM analysis. GLMMs were fitted using the *glmer* function in *lme4* (*v1.1-2* Bates et al., 2015) to establish global models for model selection of feeding guild abundances, aboveground plant biomass, and average seed set per flower. Feeding guild abundances and the number of active inflorescences were pooled across the year for each plant and plot to include both floral fitness outcomes and arthropod incidence in a single analysis. Model selection was conducted separately for each data subset. All interactions hypothesized in our original SEM between feeding guilds, clothianidin treatment, and the number of inflorescences were included in each global model. Plant species was also included as a random effect in every model. The *dredge* function from the *MuMin* package (Bartoń, 2023) was then used to identify the best model for each dependent variable in our SEM. GLMMs were then used for a piecewise SEM analysis using the *psem* function (Lefcheck, 2016). Overall model goodness of fit was assessed using the *fisherC* function (Lefcheck, 2016). Only models with a *p-*value >0.05 were considered a good fit for the data.

Multicollinearity among independent variables was assessed by calculating variance inflation factors (VIF). VIF scores from all models showed low correlation between predictor variables, indicating that our covariates were sufficiently independent. Residuals from all models were assessed visually and via Shapiro-Wilk tests of normality. Heteroskedasticity was assessed visually and via Breusch-Pagan tests for all models.

### Statistical Analysis - PERMANOVA

To assess family-level arthropod community differences, we utilized non-parametric multivariate analysis of variance (PERMANOVA) via the *adonis2* function from the *vegan* package (Oksanen J et al., 2024), with 999 permutations. Community structure comparisons were made between clothianidin and control, between plant species, and among interactions of plant species and treatment. Permutations were constrained using date as a stratum term to account for seasonal variation. For all comparisons, pairwise Bray-Curtis dissimilarity indices were used. Homogeneity of variance was assessed for all corresponding PERMANOVA analyses. Differences between clothianidin-treated and control plots and between plant species were visualized using non-metric multidimensional scaling (NMDS) ordinations.

All statistical modeling was conducted in R v4.3.2 (R Core team 2024).

## Results

Our structural equation models revealed that clothianidin soil contamination did not uniformly affect arthropod-feeding guilds or plant fitness metrics (Figures 1 & 2). Instead, the plant trait of floral inflorescence abundance was a much more influential factor on arthropod feeding guild abundance (Figures 1 & 2).

**Figure 1.**
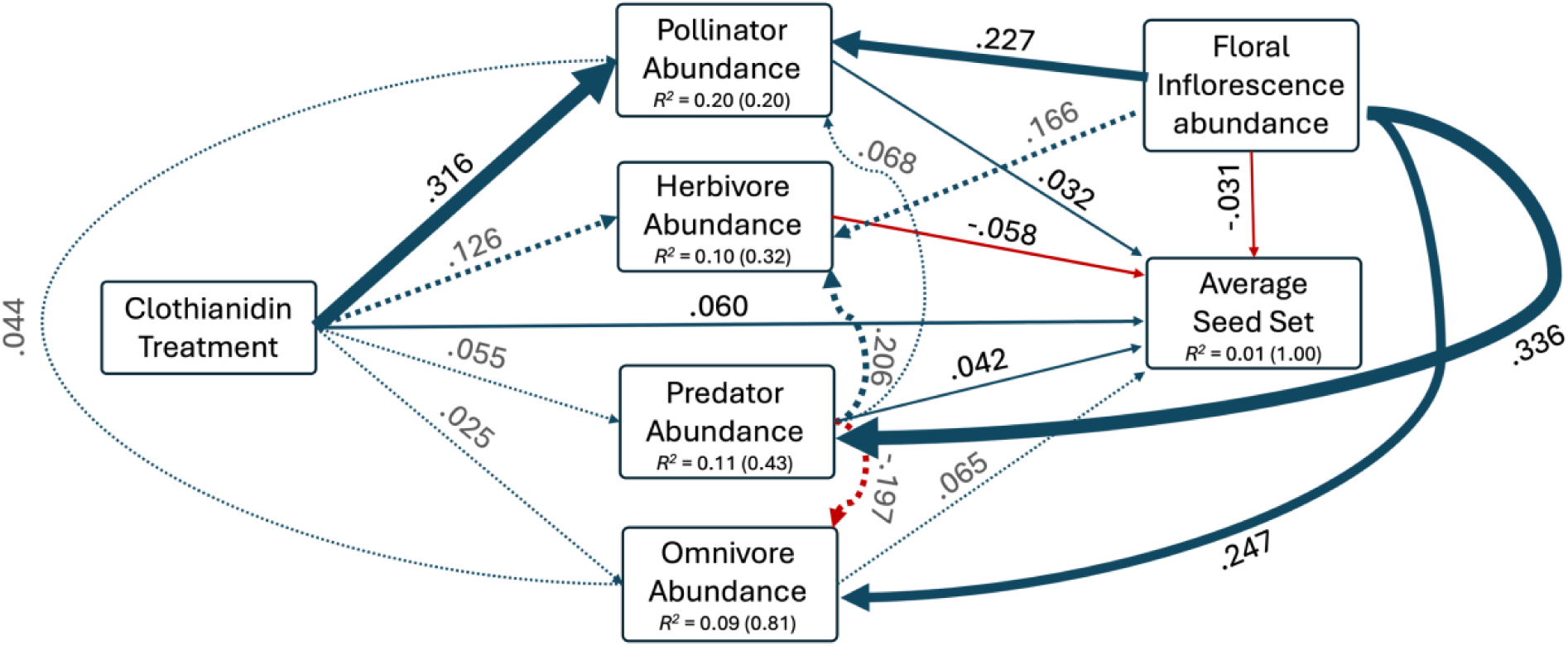
Structural equation model relating to the effects of clothianidin soil contamination, arthropod feeding guild abundances, and plant inflorescence production on average seed set per flowering head at the end of the growing season. Arrows represent directional effects between variables. Solid lines represent statistically significant relationships (*p-*value < 0.05), while dotted lines represent non-significant relationships (*p-*value > 0.05). Arrow widths are scaled by the standardized coefficients (Table 1) and are colored based on the directionality of their effect (blue = positive, red = negative). Marginal (fixed effects only) and conditional (fixed and random effects) *R^2^* values are also reported for each response variable; conditional values are in parentheses.

**Figure 2.**
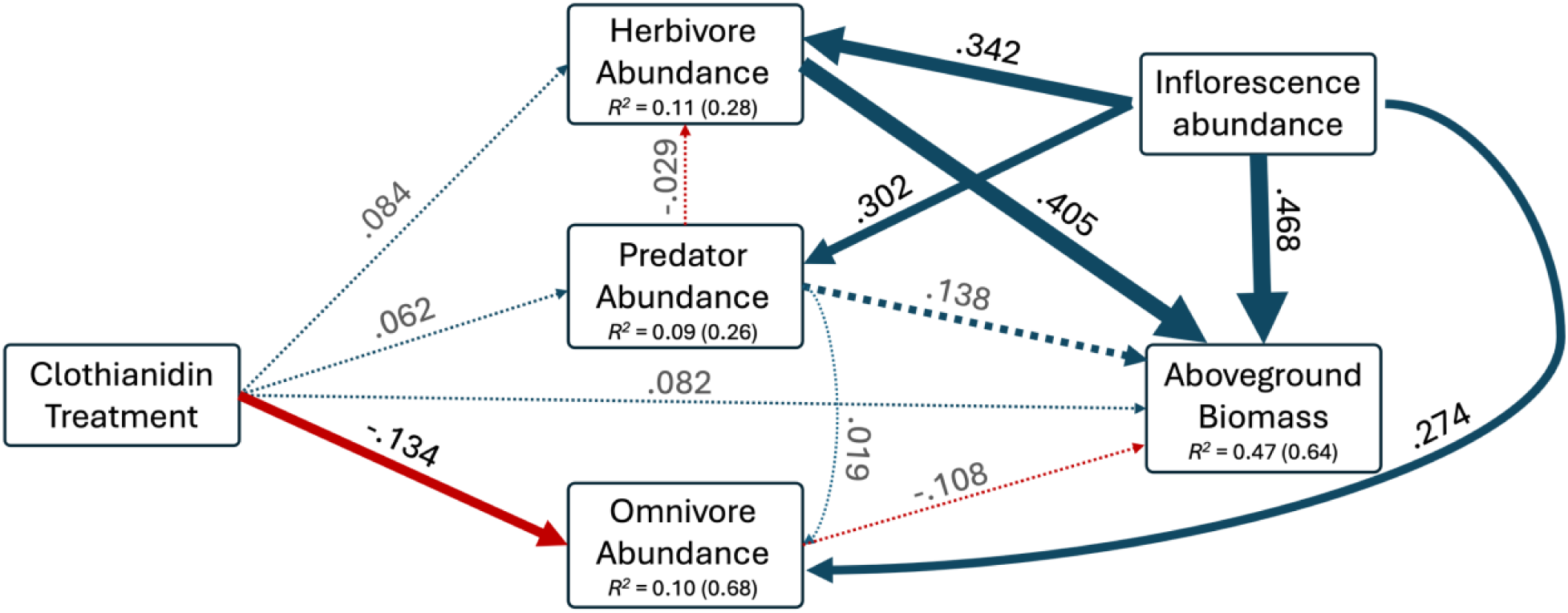
Structural equation model relating to the effects of clothianidin soil contamination, arthropod feeding guild abundances, and plant inflorescence production on average plant aboveground biomass at the end of the growing season. Arrows represent directional effects between variables. Solid lines represent statistically significant relationships (*p-*value < 0.05), while dotted lines represent non-significant relationships (*p-*value > 0.05). Arrow widths are scaled by the standardized coefficients (Table 1) and are colored based on the directionality of their effect (blue = positive, red = negative). Marginal (fixed effects only) and conditional (fixed and random effects) *R^2^* values are also reported for each response variable; conditional values are in parentheses.

### Seed set SEM

Pollinator abundance was positively associated with clothianidin contamination (*t* = 2.656, *p =* 0.008) (Table & Figure 1). Additionally, clothianidin had a positive effect on average seed set, independent from arthropod interactions (*t* = 9.373, *p =* 0.000) (Table & Figure 1).

Aside from clothianidin contamination, floral inflorescence abundance had the largest impact on feeding guild abundances. Pollinator, predator, and omnivore abundances were all positively associated with floral inflorescence abundance (*t* = 2.037, *p =* 0.042; *t* = 2.037, *p =* 0.0003; *t* = 3.384*, p =* 0.0007, respectively) (Table & Figure 1). Herbivore abundance had a positive but insignificant association with floral inflorescence abundance (*t* = 1.302, *p =* 0.193) (Table & Figure 1). Arthropod intraguild interactions had no significant causal links, however predator abundance had a marginally significant negative effect on omnivore abundance (*t* = -1.675, *p =* 0.094) (Table & Figure 1). Seed set was positively associated with pollinator abundance, predator abundance, and clothianidin contamination (*t* = 5.845, *p =* 0.000; *t* = 5.674, *p =* 0.000; *t* = 9.373, *p =* 0.000, respectively) and negatively associated with herbivore abundance and floral inflorescence abundance (*t* = -7.140, *p =* 0.000; *t* = -3.419, *p =* 0.0006) (Table & Figure 1).

In addition to the direct effects outlined above, both clothiandin contamination and floral inflorescence abundance indirectly influenced average seed set in a positive direction. Clothianidin had a positive indirect effect on average seed set, mediated by pollinator abundance (Figure 1). Floral inflorescence abundance had positive indirect effects on average seed set, mediated by pollinator and predator abundance (Figure 1).

### Aboveground biomass SEM

Omnivore abundance was the only feeding guild that was significantly influenced by clothianidin contamination (*t* = -2.007, *p =* 0.045) (Table & Figure 2). Floral inflorescence abundance was positively associated with predator abundance, omnivore abundance, herbivore abundance, and aboveground biomass (*t* = 3.325, *p =* 0.001; *t* = 3.263*, p =* 0.001; *t* = 3.182, *p =* 0.002; *t* = 3.840, *p =* 0.0003, respectively) (Table & Figure 2). In addition to direct effects, floral inflorescence abundance had a positive indirect effect on aboveground biomass, mediated by herbivore abundance (Figure 2). No significant intraguild arthropod interactions were identified. Herbivore abundance was positively associated with aboveground biomass (*t* = 3.884, *p =* 0.0002) (Table & Figure 2).

### Arthropod community structure

Arthropod community assemblages – determined from our PERMANOVA - were significantly affected by plant species (*F* = 8.584, *p* = 0.001) but were not significantly affected by clothianidin contamination (*F* = 0.823, *p* = 0.552), or the interaction between plant species and clothianidin contamination (*F* = 0.821, *p* = 0.691). Homogeneity of variance analysis showed that there were no significant differences in dispersal from the centroids for clothianidin treatment (*F* = 0.059, *p* = 0.771) (Appendix S1: Table S2). However, homogeneity of variance analysis for the plant PERMANOVA showed a non-homogenous dispersion (*F* = 7.342, *p* < 0.001) (Appendix S1: Table S2), which could have an effect on estimates and significance values. Nevertheless, the NMDS ordination (Figure 3) supports a clustering pattern that confirms the importance of plant species on arthropod community structure.

**Figure 3.**
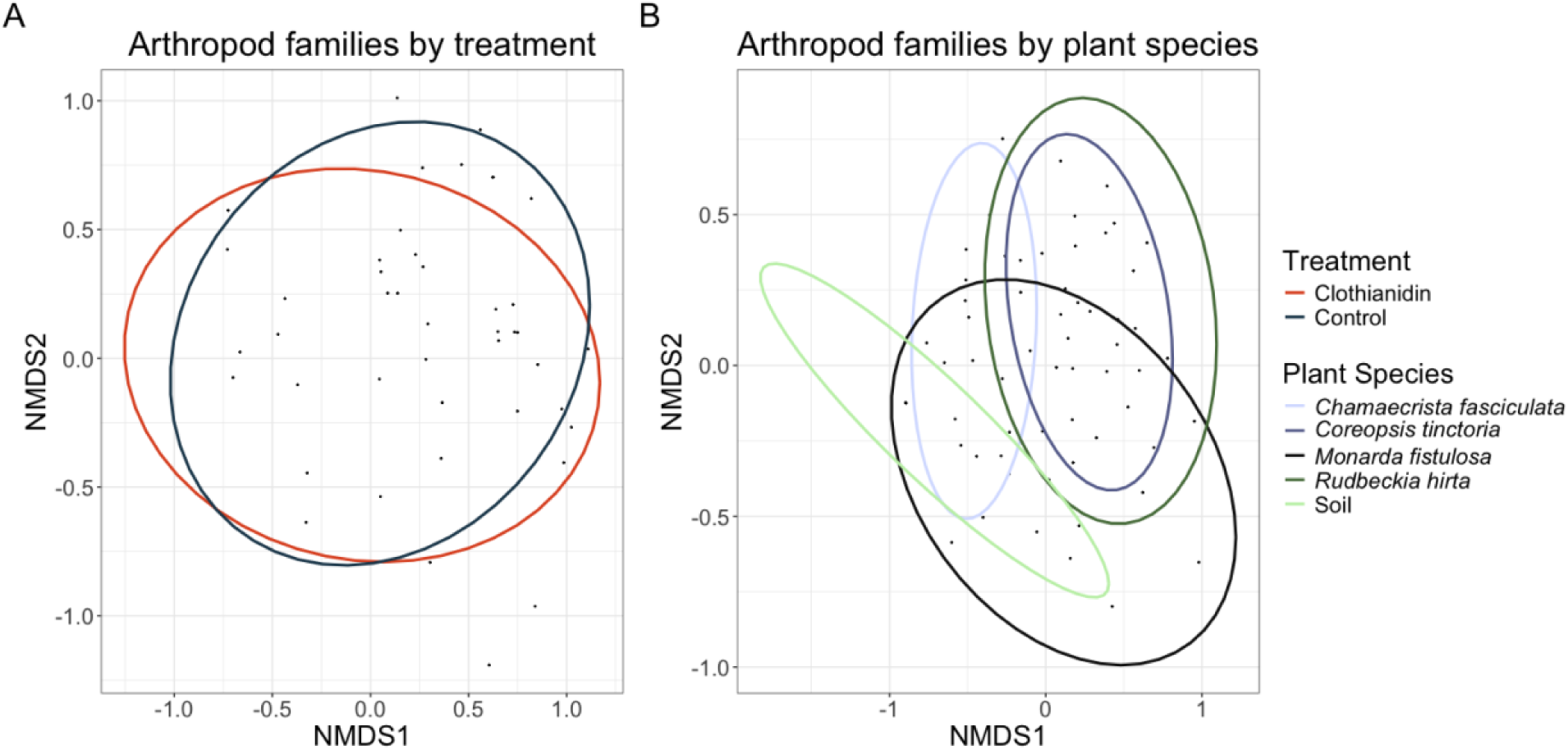
Non-metric multidimensional scaling (NMDS) ordination of arthropod family assemblages by (A) treatment and (B) plant species.

## Discussion

In this in situ manipulated field experiment in a prairie restoration ecosystem, we showed that, in general, floral characteristics rather than neonicotinoid contamination, were primarily responsible for arthropod feeding guild abundances. Similarly, floral species significantly impacted arthropod family structure, while clothianidin did not have a significant impact on arthropod family structure. However, we found that clothianidin contamination had a positive effect on the seed set of young prairie forbs by increasing seed set directly and indirectly through pollinator abundance, which also had a positive effect on seed set. This study provides a valuable framework for characterizing the relative importance of community-level interactions between ecologically important groups in an early restoration.

### Seed Set SEM outcomes

Long-term habitat stability in restoration ecosystems relies heavily on the ability of prairie plants to establish and reproduce (Benson and Hartnett, 2006). We found that clothianidin contamination in early prairie plant establishment can positively influence seed set directly and indirectly through the enhanced recruitment of pollinators. These results suggest that the ability of flowering plants to reproduce in newly established restorations is incidentally enhanced when neonicotinoids are present. Bees were the dominant pollinator observed in the study, representing over 82% of total visits. This positive association between bees and neonicotinoids corroborates earlier studies that have shown a proclivity of bees to consume contaminated resources (Kessler et al., 2015; Arce et al., 2018) and nest in contaminated areas (Tetlie and Harmon-Threatt, 2024). Despite a potential enhancement of foraging habitat caused by neonicotinoid contamination, sublethal exposure to these chemicals is likely detrimental to bee community health over time.

Exposure to neonicotinoids decreases fecundity (Sandrock et al., 2014), larval development rate (Anderson and Harmon-Threatt, 2019), adult longevity (Anderson and Harmon-Threatt, 2019) reproduction, and population growth rate (Stuligross and Williams, 2021) in non-*Apis* species. It is therefore possible that despite clothianidin contamination positively affecting pollinator recruitment to prairie flowers and subsequent floral fitness, chronic exposure to neonicotinoid-contaminated habitats could negatively impact pollinator communities over time.

While predators and herbivores were not impacted by neonicotinoid exposure, they did have significant positive and negative impacts on seed set, respectively. These findings are consistent with other studies which indicate that neonicotinoid contamination does not drastically disrupt these ecological groups, particularly at the levels found in restoration habitats (Alford and Krupke, 2017; Ding et al., 2018) (Appendix S1: Table S3). Additionally, predators were positively influenced by floral inflorescence abundance. This positive association was likely driven by arthropod predators that were commonly observed on flowers, such as crab spiders (Thomisidae), jumping spiders (Salticidae), and ambush bugs (*Phymata* sp.). These predators often prey upon presumed pollinators (Morse, 1984; Gil-Santana et al., 2015; Trindade-Santos et al., 2023) – which were also positively influenced by inflorescence abundance. However, we did not see any significant interaction between pollinators and predators in our SEM. The non-significance between predators and pollinators, in addition to the non-significant predator-herbivore interaction, is likely caused by a diluting effect of grouping flower-associated and non-flower-associated predators.

While our findings indicate that clothianidin contamination could enhance floral fitness, successful plant establishment does not guarantee the successful restoration of ecological function (Hansen and Gibson, 2014). The lack of causal links between clothianidin and certain feeding guild abundances likely does not capture potential sublethal effects that arthropods may be experiencing, which could have prolonged consequences on arthropod community structure and function and subsequent effects on the plant community.

### Aboveground biomass SEM outcomes

Aboveground biomass metrics are useful for assessing short- and long-term productivity and fitness in prairie plants (Biondini, 2007). Neonicotinoids have been shown to increase plant height in certain agricultural commodities (Preetha and Stanley, 2012), however this has not previously been examined in prairie plants. Our results show that clothianidin contamination had neither significant direct nor indirect effects on aboveground biomass of the examined prairie forbs.

Though we anticipated herbivores to decrease in abundance on contaminated plants, we found they were not significantly affected by clothianidin contamination. This finding is consistent with row crop literature (Alford and Krupke, 2017). While clothianidin levels in our plants (Appendix S1: Table S1) would be considered high in restoration habitats (Cheng, 2021), they would not be regarded as high enough to provide meaningful control in row crop agriculture (Alford and Krupke, 2017). In fact, certain crop pests, such as the black cutworm (*Agrotis ipsilon*), exhibit the hormetic effect of increasing their reproductive rate when treated with sublethal doses of clothianidin (Ding et al., 2018). Furthermore, certain folivores, such as *A. ipsilon*, are far less susceptible to clothianidin than other arthropod groups, such as bees (Costa et al., 2015; Beadle et al., 2019; Mayack and Boff, 2019). This means that herbivore communities are likely less harmed by neonicotinoid contamination and could instead be augmented by low levels of neonicotinoids like those found in prairie restorations. Additionally, neonicotinoid contamination could exhibit a selective pressure on certain herbivore groups, therefore increasing opportunities for the evolution of resistant arthropod populations (Brewer et al., 2024).

Predator abundance was similarly not significantly influenced by clothianidin contamination. Arthropod predator outcomes relative to neonicotinoid contamination are currently mixed and therefore warrant further study (Prabhaker et al., 2011, 2017; Rix and Cutler, 2020). Sublethal effects of neonicotinoids on predators range from hormetic effects on the reproduction of pentatomid predators (Rix and Cutler, 2020), causing significant decreases in overall predator biomass (Harmon et al., 2023), and significantly impairing movement and consumption in certain predatory beetles (Tooming et al., 2017). While many of these findings are taxon-specific and often lack the biological context of a field setting, the variation in outcomes could be indicative of longitudinal trends that could cause restructuring or instability within arthropod predator feeding guilds in these contaminated ecosystems. Long-term studies that maintain biological relevance are therefore required to assess prolonged trends.

While not significant in our seed set model (see above), omnivores were significantly reduced by clothianidin in our aboveground biomass model. This trend is likely driven by ant abundance, specifically those found on *C. fasciculata,* which represent 66% of all omnivores observed in this study. Ants are commonly found tending to extrafloral nectaries (EFNs) on *C. fasciculata* and often provide defense against herbivores (Fehling, 2022). In addition to a reduction in overall abundance, we did not observe any herbivore reductions as a result of ant abundance. Ants feeding on these EFNs are potentially exposed to greater amounts of neonicotinoids than other omnivores not feeding directly on nectar. The observed reduction in omnivores is consistent with previous work showing reduced foraging of ants exposed to neonicotinoids (Thiel and Köhler, 2016). Furthermore, like other eusocial hymenopterans, prolonged neonicotinoid exposure reduces queen fecundity (Wu-Smart and Spivak, 2016; Schläppi et al., 2020). The negative effects on ants could be particularly detrimental, as ants occupy numerous ecological niches and provide a multitude of ecosystem services in prairie ecosystems (Nemec, 2014; Wills and Landis, 2018).

Despite a lack of influence from neonicotinoid exposure, aboveground biomass was both directly and indirectly influenced by inflorescence abundance. The indirect effect of floral inflorescence on aboveground biomass was mediated by herbivore abundance and is in line with earlier studies that have shown an increase in herbivory associated with a corresponding increase in inflorescence size (Sletvold and Grindeland, 2008). Furthermore, herbivory can induce compensatory growth and, in some cases, overcompensation in a wide variety of plants, including the plant families examined in this study (Garcia and Eubanks, 2019). It is also likely that the positive association between inflorescence abundance and aboveground biomass is simply related to an increase in available resources associated with larger plant size (Younginger et al., 2017).

### Arthropod community structure outcomes

Consistent with findings from our SEMs highlighting the importance of plants on arthropods, results from our PERMANOVA indicate that arthropod community structure at the family level was significantly impacted by floral species but not by either clothianidin contamination or an interaction between plant species and clothianidin. Furthermore, this suggests that plant resources and subsequent potential ecological niches for arthropods vary between plant families, causing significant differences in arthropod community associations between plants. This can be seen when comparing the two plants from the Asteraceae – which had the most similar arthropod community structure (Figure 3) – with the other plants used in this study. Previous work supports the importance of plant species composition in shaping arthropod community structure (Schaffers et al., 2008) and though we added neonicotinoids to our study, we found similar results. We postulate that improving plant community composition is the most important factor in promoting resiliency and stability in the arthropod community of early restorations and degraded landscapes.

While the results from the PERMANOVA and SEM cannot be compared directly due to differences in the grouping of arthropods, both sets of analyses suggest that plant species and floral characteristics are more influential factors on arthropod community structure and feeding guild abundances than neonicotinoid contamination.

### Conclusions

Neonicotinoid contamination of new and existing natural habitats will likely continue and potentially increase for the foreseeable future as the majority of corn and soy acreage planted in the United States is now treated with some formulation of neonicotinoid (Douglas and Tooker, 2015), and there is a trend of increasing per-seed application rates (Tooker et al., 2017). Our study indicates that establishing a diverse plant community within agroecosystems is arguably more critical to the stability of arthropod community structure and feeding guild abundance than the potential negative impacts caused by neonicotinoid contamination. However, because of the potential generational impacts of sublethal neonicotinoid exposure, further longitudinal studies are required to more accurately assess the potential impacts of neonicotinoid contamination on restoration ecosystems. While not examined in this study, one potential explanation for the minimal effect of neonicotinoids within these systems could be that the insect communities found in these regions are reduced to disturbance-tolerant species after years of habitat degradation, natural area loss, and persistent pesticide usage.

**Table 1.** Estimates of regression coefficients and their standard errors (SE) from the average seed set structural equation model (Fig. 1). Standardized coefficients are shown in bold when significant (P < 0.05).

| Response | Predictor | Estimate | SE | df | t-value | p-value | Std. Estimate |
| --- | --- | --- | --- | --- | --- | --- | --- |
| *Pollinator abundance* | Clothianidin | 0.6687 | 0.2517 | 44 | 2.6564 | **0.0079** | **0.3164** |
|  | Floral inflorescences | 0.0156 | 0.0077 | 44 | 2.0372 | **0.0416** | **0.2270** |
|  | Predator abundance | 0.0320 | 0.0613 | 44 | 0.5223 | 0.6014 | 0.0681 |
|  | Omnivore abundance | 0.0040 | 0.0115 | 44 | 0.3491 | 0.7270 | 0.0438 |
| *Herbivore abundance* | Clothianidin | 0.1958 | 0.1920 | 44 | 1.0200 | 0.3077 | 0.1257 |
|  | Predator abundance | 0.0695 | 0.0447 | 44 | 1.5539 | 0.1202 | 0.2006 |
|  | Floral inflorescences | 0.0084 | 0.0065 | 44 | 1.3024 | 0.1928 | 0.1663 |
| *Omnivore abundance* | Clothianidin | 0.0775 | 0.2036 | 44 | 0.3804 | 0.7036 | 0.0251 |
|  | Predator abundance | -0.1351 | 0.0807 | 44 | -1.6751 | 0.0939 | -0.1969 |
|  | Floral inflorescences | 0.0248 | 0.0073 | 44 | 3.3838 | **0.0007** | **0.2470** |
| *Predator abundance* | Clothianidin | 0.1054 | 0.2073 | 44 | 0.5083 | 0.6112 | 0.0553 |
|  | Floral inflorescences | 0.0208 | 0.0058 | 44 | 3.6131 | **0.0003** | **0.3356** |
| *Average seed set* | Pollinator abundance | 0.0345 | 0.0059 | 44 | 5.8449 | **0.0000** | **0.0322** |
|  | Herbivore abundance | -0.0290 | 0.0041 | 44 | -7.1404 | **0.0000** | **-0.0584** |
|  | Clothianidin | 0.2044 | 0.0218 | 44 | 9.3732 | **0.0000** | **0.0596** |
|  | Omnivore abundance | 0.0096 | 0.0056 | 44 | 1.7054 | 0.0881 | 0.0649 |
|  | Predator abundance | 0.0319 | 0.0056 | 44 | 5.6741 | **0.0000** | **0.0418** |
|  | Floral inflorescences | -0.0035 | 0.0010 | 44 | -3.4185 | **0.0006** | **-0.0314** |

**Table 2.** Estimates of regression coefficients and their standard errors (SE) from the aboveground biomass structural equation model (Fig. 2). Standardized coefficients are shown in bold when significant (P < 0.05).

| Response | Predictor | Estimate | SE | df | t-value | p-value | Std. Estimate |
| --- | --- | --- | --- | --- | --- | --- | --- |
| *Omnivore abundance* | Clothianidin | -0.3565 | 0.1777 | 80 | -2.0067 | **0.0448** | **-0.1343** |
|  | Predator abundance | 0.0135 | 0.0580 | 80 | 0.2335 | 0.8153 | 0.0192 |
|  | Floral inflorescences | 0.0237 | 0.0073 | 80 | 3.2625 | **0.0011** | **0.2741** |
| *Herbivore abundance* | Clothianidin | 0.1481 | 0.1594 | 80 | 0.9289 | 0.3530 | 0.0842 |
|  | Predator abundance | -0.0135 | 0.0475 | 80 | -0.2838 | 0.7765 | -0.0289 |
|  | Floral inflorescences | 0.0196 | 0.0062 | 80 | 3.1820 | **0.0015** | **0.3419** |
| *Predator abundance* | Clothianidin | 0.1649 | 0.2252 | 80 | 0.7324 | 0.4639 | 0.0615 |
|  | Floral inflorescences | 0.0263 | 0.0079 | 80 | 3.3253 | **0.0009** | **0.3015** |
| *Aboveground biomass (g)* | Clothianidin | 0.8980 | 0.9038 | 80 | 0.9935 | 0.3238 | 0.0820 |
|  | Herbivore abundance | 0.7157 | 0.1843 | 80 | 3.8842 | **0.0002** | **0.4046** |
|  | Floral inflorescences | 0.1668 | 0.0434 | 80 | 3.8404 | **0.0003** | **0.4677** |
|  | Predator abundance | 0.4012 | 0.2953 | 80 | 1.3588 | 0.1784 | 0.1381 |
|  | Omnivore abundance | -0.0622 | 0.0898 | 80 | 0.4941 | 0.4941 | -0.1084 |

**Table 3.** Estimates, goodness of fit, and *p-*values for PERMANOVA analyses (Fig. 3).

| | *df* | Sum of squares | $R^2$ | *f*-value | *p*-value |
| --- | --- | --- | --- | --- | --- |
| *Clothianidin* | 1 | 0.156 | 0.009 | 0.823 | 0.552 |
| *Plant species* | 4 | 6.505 | 0.384 | 8.584 | **0.001** |
| *Clothianidin x plant species* | 4 | 0.622 | 0.037 | 0.821 | 0.691 |

## Supporting information

Supplemental Tables 1-3 and supplemental Figure 1, and will be used for the link to the file on the preprint site.

## 4) Acknowledgments

The authors would like to thank Adam Dolezal, May Berenbaum, and Anthony Yannarell for their input in the establishment of this project, Jamie Ellis and Nate Hudson for their assistance in site scouting and establishment of the sites used in this project, and Nick Anderson, Annaliese Wargin, Virginia Roberts, Kristine Schoenecker, Jamilyn Martin, and Vi Aldunate for providing feedback on the manuscript.

The authors would also like to thank the following funding sources: This research was supported by USDA-NIFA, and the School of Integrative Biology at the University of Illinois at Urbana-Champaign (Fred H. Schmidt Summer Scholars Award, Francis M. and Harlie M. Clark Research Support Grant, and Harley J. Van Cleave Research Award).

## 5) Author Contributions

JT: Conceptualization, Data curation, Formal analysis, Funding acquisition, Investigation, Methodology, Project administration, Resources, Software, Validation, Visualization, Writing – original draft, Writing – review & editing. AH: Conceptualization, Funding acquisition, Project administration, Resources, Supervision, Validation, Writing – review & editing.

## Boxes

N/A

## References

Alford, A., and Krupke, C. H. (2017). Translocation of the neonicotinoid seed treatment clothianidin in maize. PLoS One 12, 1–19. doi: 10.1371/journal.pone.0173836.

Anderson, N. L., and Harmon-Threatt, A. N. (2019). Chronic contact with realistic soil concentrations of imidacloprid affects the mass , immature development speed , and adult longevity of solitary bees. Sci. Rep., 1–9. doi: 10.1038/s41598-019-40031-9.

Angelella, G. M., and O’Rourke, M. E. (2017). Pollinator habitat establishment after organic and no-till seedbed preparation methods. HortScience 52, 1349–1355. doi: 10.21273/HORTSCI11962-17.

Arce, A. N., Rodrigues, A. R., Yu, J., Colgan, T. J., Wurm, Y., and Gill, R. J. (2018). Foraging bumblebees acquire a preference for neonicotinoid-treated food with prolonged exposure. Proc. R. Soc. B Biol. Sci. 285, 8–11. doi: 10.1098/rspb.2018.0655.

Aseperi, A. K., Busquets, R., Hooda, P. S., Cheung, P. C. W., and Barker, J. (2020). Behaviour of neonicotinoids in contrasting soils. J. Environ. Manage. 276, 111329. doi: 10.1016/j.jenvman.2020.111329.

Badua, S. A., Sharda, A., Strasser, R., and Ciampitti, I. (2021). Ground speed and planter downforce influence on corn seed spacing and depth. Precis. Agric. 22, 1154–1170. doi: 10.1007/s11119-020-09775-7.

Bartoń, K. (2023). MuMIn: Multimodal Inference. R Packag. ver. 1.47.5.

Bates, D., Mächler, M., Bolker, B. M., and Walker, S. C. (2015). Fitting linear mixed-effects models using lme4. J. Stat. Softw. 67, 1–48. doi: 10.18637/jss.v067.i01.

Beadle, K., Singh, K. S., Troczka, B. J., Randall, E., Zaworra, M., Zimmer, C. T., et al. (2019). Genomic insights into neonicotinoid sensitivity in the solitary bee Osmia bicornis. PLoS Genet. 15, 1–19. doi: 10.1371/journal.pgen.1007903.

Benson, E. J., and Hartnett, D. C. (2006). The role of seed and vegetative reproduction in plant recruitment and demography in tallgrass prairie. Plant Ecol. 187, 163–178. doi: 10.1007/s11258-005-0975-y.

Biondini, M. (2007). Plant diversity, production, stability, and susceptibility to invasion in restored northern tall grass prairies (United States). Restor. Ecol. 15, 77–87. doi: 10.1111/j.1526-100X.2006.00192.x.

Borror, D. J., White, R. E., Society, N. A., Federation, N. W., and Institute, R. T. P. (1998). A Field Guide to Insects: America North of Mexico. Houghton Mifflin Available at: https://books.google.com/books?id=yXK2QgAACAAJ.

Brewer, M. J., Umina, P. A., and Elliott, N. C. (2024). Pest Management for Spatially Variable Arthropod Pests in Large-scale Agroecosystems. CABI Books, 27–43. doi: 10.1079/9781800622777.0002.

Burghardt, K. T., and Tallamy, D. W. (2013). Plant origin asymmetrically impacts feeding guilds and life stages driving community structure of herbivorous arthropods. Divers. Distrib. 19, 1553–1565. doi: 10.1111/ddi.12122.

Cheng, S.-H. (2021). Quantification of neonicotinoid residues in soils and dust drift in conservation reserve program fields in Illinois, USA.

Costa, L. M., Grella, T. C., Barbosa, R. A., Malaspina, O., and Nocelli, R. C. F. (2015). Determination of acute lethal doses (LD50 and LC50) of imidacloprid for the native bee Melipona scutellaris Latreille, 1811 (Hymenoptera: Apidae). Sociobiology 62, 578–582. doi: 10.13102/sociobiology.v62i4.792.

Ding, J., Zhao, Y., Zhang, Z., Xu, C., and Mu, W. (2018). Sublethal and Hormesis Effects of Clothianidin on the Black Cutworm (Lepidoptera: Noctuidae). J. Econ. Entomol. 111, 2809–2816. doi: 10.1093/jee/toy254.

Disque, H. H., Hamby, K. A., Dubey, A., Taylor, C., and Dively, G. P. (2019). Effects of clothianidin-treated seed on the arthropod community in a mid-Atlantic no-till corn agroecosystem. Pest Manag. Sci. 75, 969–978. doi: 10.1002/ps.5201.

Douglas, M. R., and Tooker, J. F. (2015). Large-scale deployment of seed treatments has driven rapid increase in use of neonicotinoid insecticides and preemptive pest management in U.S. Field crops. Environ. Sci. Technol. 49, 5088–5097. doi: 10.1021/es506141g.

Dudley, N., and Alexander, S. (2017). Agriculture and biodiversity: a review. Biodiversity 18, 45–49. doi: 10.1080/14888386.2017.1351892.

Fehling, L. S. (2022). Reward Complementarity and Context Dependency in Multispecies Mutualist Interactions in Partridge Pea (Chamaecrista fasciculata). Available at: http://rave.ohiolink.edu/etdc/view?acc_num=miami1653493649405494.

Garcia, L. C., and Eubanks, M. D. (2019). Overcompensation for insect herbivory: a review and meta-analysis of the evidence. Ecology 100, 1–14. doi: 10.1002/ecy.2585.

Gil-Santana, H. R., Forero, D., and Weirauch, C. (2015). “Assassin Bugs (Reduviidae Excluding Triatominae) BT - True Bugs (Heteroptera) of the Neotropics,” in, eds. A. R. Panizzi and J. Grazia (Dordrecht: Springer Netherlands), 307–351. doi: 10.1007/978-94-017-9861-7_12.

Goulson, D. (2013). An overview of the environmental risks posed by neonicotinoid insecticides. J. Appl. Ecol. 50, 977–987. doi: 10.1111/1365-2664.12111.

Grman, E., Lau, J. A., Schoolmaster, D. R., and Gross, K. L. (2010). Mechanisms contributing to stability in ecosystem function depend on the environmental context. Ecol. Lett. 13, 1400– 1410. doi: 10.1111/j.1461-0248.2010.01533.x.

Hall, M. J., Zhang, G., O’Neal, M. E., Bradbury, S. P., and Coats, J. R. (2022). Quantifying neonicotinoid insecticide residues in milkweed and other forbs sampled from prairie strips established in maize and soybean fields. Agric. Ecosyst. Environ. 325, 107723. doi: 10.1016/j.agee.2021.107723.

Hansen, M. J., and Gibson, D. J. (2014). Use of multiple criteria in an ecological assessment of a prairie restoration chronosequence. Appl. Veg. Sci. 17, 63–73. doi: 10.1111/avsc.12051.

Harmon, G. T., Harmon-Threatt, A. N., and Anderson, N. L. (2023). Changes in predator biomass may mask the negative effects of neonicotinoids on primary consumers in field settings. Insect Conserv. Divers. 16, 298–305. doi: 10.1111/icad.12625.

Harvey, J. A., van der Putten, W. H., Turin, H., Wagenaar, R., and Bezemer, T. M. (2008). Effects of changes in plant species richness and community traits on carabid assemblages and feeding guilds. Agric. Ecosyst. Environ. 127, 100–106. doi: 10.1016/j.agee.2008.03.006.

Henrik Barmentlo, S., Schrama, M., De Snoo, G. R., Van Bodegom, P. M., Van Nieuwenhuijzen, A., and Vijver, M. G. (2021). Experimental evidence for neonicotinoid driven decline in aquatic emerging insects. Proc. Natl. Acad. Sci. U. S. A. 118, 2–9. doi: 10.1073/pnas.2105692118.

Hladik, M. L., Main, A. R., and Goulson, D. (2018). Environmental Risks and Challenges Associated with Neonicotinoid Insecticides. Environ. Sci. Technol. 52, 3329–3335. doi: 10.1021/acs.est.7b06388.

Humann-Guilleminot, S., Binkowski, Ł. J., Jenni, L., Hilke, G., Glauser, G., and Helfenstein, F. (2019). A nation-wide survey of neonicotinoid insecticides in agricultural land with implications for agri-environment schemes. J. Appl. Ecol. 56, 1502–1514. doi: 10.1111/1365-2664.13392.

Kessler, S. C., Tiedeken, E. J., Simcock, K. L., Derveau, S., Mitchell, J., Softley, S., et al. (2015). Bees prefer foods containing neonicotinoid pesticides. Nature 521, 74–76. doi: 10.1038/nature14414.

Khan, S. U. (2016). Pesticides in the soil environment. Elsevier.

Kurtz, C. (2013). A Practical Guide to Prairie Reconstruction: Second *Edition*. University of Iowa Press Available at: https://books.google.com/books?id=tIKpxcyB2DYC.

Lefcheck, J. S. (2016). piecewiseSEM: Piecewise structural equation modelling in r for ecology, evolution, and systematics. Methods Ecol. Evol. 7, 573–579. doi: 10.1111/2041-210X.12512.

Longcore, T. (2003). Terrestrial arthropods as indicators of ecological restoration success in coastal sage scrub (California, U.S.A.). Restor. Ecol. 11, 397–409. doi: 10.1046/j.1526-100X.2003.rec0221.x.

Main, A. R., Headley, J. V., Peru, K. M., Michel, N. L., Cessna, A. J., and Morrissey, C. A. (2014). Widespread use and frequent detection of neonicotinoid insecticides in wetlands of Canada’s prairie pothole region. PLoS One 9. doi: 10.1371/journal.pone.0092821.

Main, A. R., Webb, E. B., Goyne, K. W., and Mengel, D. (2018). Neonicotinoid insecticides negatively affect performance measures of non-target terrestrial arthropods: a meta-analysis. Ecol. Appl. 28, 1232–1244.

Mayack, C., and Boff, S. (2019). LD50 values may be misleading predictors of neonicotinoid toxicity across different bee species. Uludag Aricilik Derg. 19, 19–33. doi: 10.31467/uluaricilik.568251.

Morales-Rodriguez, A., and Peck, D. C. (2009). Synergies between biological and neonicotinoid insecticides for the curative control of the white grubs Amphimallon majale and Popillia japonica. Biol. Control 51, 169–180. doi: 10.1016/j.biocontrol.2009.06.008.

Morse, D. H. (1984). How Crab Spiders ( Araneae , Thomisidae ) Hunt at Flowers Author ( s ): Douglass H. Morse Published by : American Arachnological Society Stable URL : http://www.jstor.com/stable/3705362 REFERENCES Linked references are available on JSTOR for this article. 12, 307–316.

Nauen, R., and Elbert, A. (1997). Apparent tolerance of a field-collected strain of Myzus nicotianae to imidacloprid due to strong antifeeding responses. Pestic. Sci. 49, 252–258. doi: 10.1002/(SICI)1096-9063(199703)49:3<252::AID-PS521>3.0.CO;2-2.

Nemec, K. T. (2014). Tallgrass prairie ants: Their species composition, ecological roles, and response to management. J. Insect Conserv. 18, 509–521. doi: 10.1007/s10841-014-9656-2.

Nemec, K. T., Allen, C. R., Danielson, S. D., and Helzer, C. J. (2014). Responses of predatory invertebrates to seeding density and plant species richness in experimental tallgrass prairie restorations. Agric. Ecosyst. Environ. 183, 11–20. doi: 10.1016/j.agee.2013.10.024.

Oksanen J, Simpson G, Blanchet F, Kindt R, Legendre P, Minchin P, et al. (2024). Community Ecology Package - Vegan. May 21. Available at: https://github.com/vegandevs/vegan.

Pisa, L., Goulson, D., Yang, E. C., Gibbons, D., Sánchez-Bayo, F., Mitchell, E., et al. (2021). An update of the Worldwide Integrated Assessment (WIA) on systemic insecticides. Part 2: impacts on organisms and ecosystems. Environ. Sci. Pollut. Res. 28, 11749–11797. doi: 10.1007/s11356-017-0341-3.

Prabhaker, N., Castle, S. J., Naranjo, S. E., Toscano, N. C., and Morse, J. G. (2011). Compatibility of two systemic neonicotinoids, imidacloprid and thiamethoxam, with various natural enemies of agricultural pests. J. Econ. Entomol. 104, 773–781. doi: 10.1603/EC10362.

Prabhaker, N., Naranjo, S., Perring, T., and Castle, S. (2017). Comparative Toxicities of Newer and Conventional Insecticides: Against Four Generalist Predator Species. J. Econ. Entomol. 110, 2630–2636. doi: 10.1093/jee/tox202.

Preetha, G., and Stanley, J. (2012). Influence of neonicotinoid insecticides on the plant growth attributes of cotton and okra. J. Plant Nutr. 35, 1234–1245. doi: 10.1080/01904167.2012.676134.

Rexrode, M., Barrett, M., Ellis, J., Patrick, G., Vaughan, A., Felkel, J., et al. (2003). EFED risk assessment for the seed treatment of clothianidin 600FS on corn and canola. 1–91.

Rix, R. R., and Cutler, G. C. (2020). Low Doses of a Neonicotinoid Stimulate Reproduction in a Beneficial Predatory Insect. J. Econ. Entomol. 113, 2179–2186. doi: 10.1093/jee/toaa169.

Ruiz-Jaen, M. C., and Aide, T. M. (2005). Restoration success: How is it being measured? Restor. Ecol. 13, 569–577. doi: 10.1111/j.1526-100X.2005.00072.x.

Sandrock, C., Tanadini, L. G., Pettis, J. S., Biesmeijer, J. C., Potts, S. G., and Neumann, P. (2014). Sublethal neonicotinoid insecticide exposure reduces solitary bee reproductive success. Agric. For. Entomol. 16, 119–128. doi: 10.1111/afe.12041.

Schaffers, A. P., Raemakers, I. P., Sykora, K. V., and Braak, C. J. F. Ter (2008). Arthropod Assemblages Are Best Predicted By Plant. Ecology 89, 782–794.

Schläppi, D., Kettler, N., Straub, L., Glauser, G., and Neumann, P. (2020). Long-term effects of neonicotinoid insecticides on ants. Commun. Biol. 3, 1–9. doi: 10.1038/s42003-020-1066-2.

Schuldt, A., Baruffol, M., Bruelheide, H., Chen, S., Chi, X., Wall, M., et al. (2014). Woody plant phylogenetic diversity mediates bottom-up control of arthropod biomass in species-rich forests. Oecologia 176, 171–182. doi: 10.1007/s00442-014-3006-7.

Shipley, B. (2009). Confirmatory path analysis in a generalized multilevel context. Ecology 90, 363–368. doi: 10.1890/08-1034.1.

Sletvold, N., and Grindeland, J. M. (2008). Floral herbivory increases with inflorescence size and local plant density in Digitalis purpurea. Acta Oecologica 34, 21–25. doi: 10.1016/j.actao.2008.03.002.

Stuligross, C., and Williams, N. M. (2021). Past insecticide exposure reduces bee reproduction and population growth rate. Proc. Natl. Acad. Sci. U. S. A. 118, 1–6. doi: 10.1073/pnas.2109909118.

Tetlie, J., and Harmon-threatt, A. (2024). Neonicotinoid contamination in conservation areas affects bees more sharply than beetles. Front. Ecol. Evol., 1–11. doi: 10.3389/fevo.2024.1347526.

Thiel, S., and Köhler, H. R. (2016). A sublethal imidacloprid concentration alters foraging and competition behaviour of ants. Ecotoxicology 25, 814–823. doi: 10.1007/s10646-016-1638-6.

Thomas, M. C., Skelley, P. E., and Frank, J. H. (2002). American Beetles, Volume II: Polyphaga: Scarabaeoidea through Curculionoidea. 1st ed. Boca Raton: CRC Press Available at: https://books.google.com/books?id=YiPNBQAAQBAJ.

Tilman, D., Lehman, C. L., and Bristow, C. E. (1998). Diversity-stability relationships: Statistical inevitability or ecological consequence? Am. Nat. 151, 277–282. doi: 10.1086/286118.

Tooker, J. F., Douglas, M. R., and Krupke, C. H. (2017). Neonicotinoid Seed Treatments: Limitations and Compatibility with Integrated Pest Management. Agric. Environ. Lett. 2, 1–5. doi: 10.2134/ael2017.08.0026.

Tooming, E., Merivee, E., Must, A., Merivee, M. I., Sibul, I., Nurme, K., et al. (2017). Behavioural effects of the neonicotinoid insecticide thiamethoxam on the predatory insect Platynus assimilis. Ecotoxicology 26, 902–913. doi: 10.1007/s10646-017-1820-5.

Trindade-Santos, M. E., Lima Ramos, R., Silva Melo, T., Brescovit, A. D., and de Oliveira, F. F. (2023). Exotic and predatory: a spider (Araneae: Salticidae) that preys on native stingless bees (Hymenoptera: Meliponini) in Brazil. Rev. Chil. Entomol. 49, 701–706. doi: 10.35249/rche.49.4.23.04.

van der Sluijs, J. P. (2020). Insect decline, an emerging global environmental risk. Curr. Opin. Environ. Sustain. 46, 39–42. doi: 10.1016/j.cosust.2020.08.012.

Wagner, D. L., Grames, E. M., Forister, M. L., Berenbaum, M. R., and Stopak, D. (2021). Insect decline in the Anthropocene: Death by a thousand cuts. Proc. Natl. Acad. Sci. U. S. A. 118, 1–10. doi: 10.1073/PNAS.2023989118.

Whitfield, J. B., Doyen, J. T., Purcell, A. H., and Purcell, A. H. (2013). Daly and Doyen’s Introduction to Insect Biology and Diversity. Oxford University Press Available at: https://books.google.com/books?id=gwWwpwAACAAJ.

Wills, B. D., and Landis, D. A. (2018). The role of ants in north temperate grasslands: a review. Oecologia 186, 323–338. doi: 10.1007/s00442-017-4007-0.

Wu-Smart, J., and Spivak, M. (2016). Sub-lethal effects of dietary neonicotinoid insecticide exposure on honey bee queen fecundity and colony development. Sci. Rep. 6, 1–11. doi: 10.1038/srep32108.

Younginger, B. S., Sirová, D., Cruzan, M. B., and Ballhorn, D. J. (2017). Is biomass a reliable estimate of plant fitness? Appl. Plant Sci. 5, 1–8. doi: 10.3732/apps.1600094.

