## Supplemental Tables 1-3 and supplemental Figure 1, and will be used for the link to the file on the preprint site. for "Plant species and floral traits shape arthropod communities in restored prairies more than neonicotinoid contamination"

**Supporting Information (Appendix)**

**Table S1:** Feeding guild abundance and plant fitness metrics for all sampled plants within each sample pot. Plants and pots that either experienced deer damage or plant mortality were excluded from the dataset.

| ***Plot #*** | ***Treatment*** | ***Plant*** | ***Herbivore Abundance*** | ***Omnivore Abundance*** | ***Pollinator Abundance*** | ***Predator Abundance*** | ***Average Seed Set*** | ***Aboveground Biomass (g)*** |
|---|---|---|---|---|---|---|---|---|
| *1* | Clothianidin | *Chamaecrista fasciculata* | 3 | 28 | 2 | 2 | 15 | 2.1 |
| *1* | Clothianidin | *Coreopsis tinctoria* | 3 | 2 | 3 | 1 | NA | 7.88 |
| *1* | Clothianidin | *Monarda fistulosa* | 0 | 3 | 0 | 1 | NA | 6.01 |
| *1* | Clothianidin | *Rudbeckia hirta* | 8 | 3 | 1 | 3 | 295.75 | 18.15 |
| *2* | Control | *Chamaecrista fasciculata* | 3 | 16 | 2 | 3 | NA | 4.72 |
| *2* | Control | *Coreopsis tinctoria* | 6 | 3 | 4 | 3 | 53 | 13.89 |
| *2* | Control | *Monarda fistulosa* | 1 | 1 | 0 | 0 | NA | 15.12 |
| *2* | Control | *Rudbeckia hirta* | 4 | 1 | 0 | 1 | 598.33 | 8.61 |
| *3* | Clothianidin | *Chamaecrista fasciculata* | 0 | 9 | 1 | 2 | 4 | 7.44 |
| *3* | Clothianidin | *Coreopsis tinctoria* | 7 | 14 | 3 | 1 | 98.67 | 18.5 |
| *3* | Clothianidin | *Monarda fistulosa* | 0 | 1 | 0 | 1 | NA | 10.28 |
| *3* | Clothianidin | *Rudbeckia hirta* | 8 | 1 | 7 | 4 | 630 | 25.2 |
| *4* | Control | *Chamaecrista fasciculata* | 0 | 25 | 0 | 0 | 22 | 5.98 |
| *4* | Control | *Coreopsis tinctoria* | 5 | 5 | 1 | 10 | NA | 11.46 |
| *4* | Control | *Monarda fistulosa* | 0 | 3 | 0 | 0 | NA | 11.51 |
| *4* | Control | *Rudbeckia hirta* | 3 | 4 | 3 | 6 | 645.33 | 10.26 |
| *5* | Clothianidin | *Chamaecrista fasciculata* | 4 | 36 | 1 | 1 | 22 | 5.98 |
| *5* | Clothianidin | *Coreopsis tinctoria* | 4 | 2 | 1 | 5 | NA | 11.46 |
| *5* | Clothianidin | *Monarda fistulosa* | 0 | 4 | 0 | 0 | NA | 11.51 |
| *5* | Clothianidin | *Rudbeckia hirta* | 3 | 3 | 6 | 3 | 645.33 | 10.26 |
| *6* | Control | *Chamaecrista fasciculata* | 3 | 22 | 2 | 1 | 4 | 10.49 |
| *6* | Control | *Coreopsis tinctoria* | 0 | 0 | 0 | 0 | NA | 1.75 |
| *6* | Control | *Monarda fistulosa* | 4 | 4 | 0 | 0 | NA | 22.77 |

| | | | | | | | | |
|---|---|---|---|---|---|---|---|---|
| *6* | Control | *Rudbeckia hirta* | 0 | 3 | 3 | 5 | NA | 9.74 |
| *7* | Clothianidin | *Chamaecrista fasciculata* | 0 | 26 | 4 | 0 | 8 | 8.61 |
| *7* | Clothianidin | *Coreopsis tinctoria* | 8 | 5 | 4 | 1 | 173 | 19.14 |
| *7* | Clothianidin | *Monarda fistulosa* | 3 | 0 | 0 | 0 | NA | 6.84 |
| *7* | Clothianidin | *Rudbeckia hirta* | 5 | 2 | 3 | 4 | 645 | 9.22 |
| *8* | Control | *Chamaecrista fasciculata* | 5 | 18 | 1 | 0 | NA | 11.52 |
| *8* | Control | *Coreopsis tinctoria* | 6 | 2 | 0 | 0 | 126 | 11.89 |
| *8* | Control | *Monarda fistulosa* | 0 | 5 | 0 | 0 | NA | 11.43 |
| *8* | Control | *Rudbeckia hirta* | 2 | 2 | 0 | 3 | NA | 9.11 |
| *9* | Clothianidin | *Chamaecrista fasciculata* | 2 | 6 | 0 | 0 | NA | 2.97 |
| *9* | Clothianidin | *Coreopsis tinctoria* | 2 | 1 | 0 | 2 | 111.67 | 11.57 |
| *9* | Clothianidin | *Monarda fistulosa* | 0 | 0 | 0 | 0 | NA | 8.18 |
| *9* | Clothianidin | *Rudbeckia hirta* | 4 | 0 | 0 | 5 | 416.33 | 16.9 |
| *10* | Control | *Chamaecrista fasciculata* | 3 | 34 | 1 | 1 | 0.67 | 5.91 |
| *10* | Control | *Coreopsis tinctoria* | 2 | 3 | 2 | 1 | NA | 8.79 |
| *10* | Control | *Monarda fistulosa* | 1 | 1 | 0 | 0 | NA | 16.05 |
| *10* | Control | *Rudbeckia hirta* | 4 | 1 | 1 | 4 | NA | 8.59 |
| *11* | Clothianidin | *Coreopsis tinctoria* | 8 | 2 | 0 | 4 | 104 | 19.77 |
| *11* | Clothianidin | *Monarda fistulosa* | 3 | 1 | 0 | 1 | NA | 14.67 |
| *11* | Clothianidin | *Rudbeckia hirta* | 7 | 2 | 1 | 0 | NA | 12.56 |
| *12* | Control | *Chamaecrista fasciculata* | 2 | 21 | 1 | 1 | 1 | 13.05 |
| *12* | Control | *Coreopsis tinctoria* | 3 | 2 | 0 | 1 | 62.67 | 8.54 |
| *12* | Control | *Monarda fistulosa* | 1 | 3 | 0 | 0 | NA | 9 |
| *12* | Control | *Rudbeckia hirta* | 0 | 0 | 0 | 0 | 352.33 | 7.51 |
| *13* | Clothianidin | *Chamaecrista fasciculata* | 3 | 34 | 4 | 2 | 8.33 | 16.52 |
| *13* | Clothianidin | *Coreopsis tinctoria* | 7 | 0 | 3 | 0 | 81.67 | 11.25 |
| *13* | Clothianidin | *Monarda fistulosa* | 1 | 0 | 0 | 0 | NA | 8.21 |
| *13* | Clothianidin | *Rudbeckia hirta* | 1 | 1 | 1 | 2 | 705.33 | 6.95 |

| | | | | | | | | |
|---|---|---|---|---|---|---|---|---|
| *14* | Control | *Chamaecrista fasciculata* | 2 | 22 | 0 | 0 | 10 | 3.22 |
| *14* | Control | *Coreopsis tinctoria* | 7 | 7 | 2 | 4 | 68 | 20.27 |
| *14* | Control | *Monarda fistulosa* | 2 | 1 | 0 | 0 | NA | 4.69 |
| *14* | Control | *Rudbeckia hirta* | 2 | 0 | 1 | 0 | NA | 1.06 |
| *15* | Clothianidin | *Chamaecrista fasciculata* | 2 | 14 | 3 | 0 | 13 | 11.45 |
| *15* | Clothianidin | *Coreopsis tinctoria* | 7 | 1 | 2 | 0 | NA | 8.13 |
| *15* | Clothianidin | *Monarda fistulosa* | 0 | 0 | 0 | 0 | NA | 4.54 |
| *15* | Clothianidin | *Rudbeckia hirta* | 2 | 0 | 1 | 4 | 573.67 | 17.04 |
| *18* | Control | *Chamaecrista fasciculata* | 3 | 26 | 1 | 0 | 15 | 8.54 |
| *18* | Control | *Coreopsis tinctoria* | 12 | 9 | 0 | 2 | NA | 12.16 |
| *18* | Control | *Monarda fistulosa* | 3 | 3 | 0 | 1 | NA | 10.59 |
| *18* | Control | *Rudbeckia hirta* | 1 | 0 | 0 | 0 | NA | 3.85 |
| *20* | Control | *Chamaecrista fasciculata* | 2 | 30 | 1 | 1 | 8 | 4.53 |
| *20* | Control | *Coreopsis tinctoria* | 3 | 4 | 3 | 0 | NA | 4.61 |
| *20* | Control | *Monarda fistulosa* | 2 | 10 | 0 | 1 | NA | 11.87 |
| *20* | Control | *Rudbeckia hirta* | 5 | 1 | 2 | 0 | 298 | 4.58 |
| *24* | Control | *Chamaecrista fasciculata* | 4 | 20 | 0 | 2 | NA | 12.88 |
| *24* | Control | *Coreopsis tinctoria* | 8 | 4 | 1 | 2 | 144.67 | 19.68 |
| *24* | Control | *Monarda fistulosa* | 4 | 0 | 0 | 1 | NA | 11.44 |
| *24* | Control | *Rudbeckia hirta* | 1 | 0 | 0 | 2 | NA | 4.19 |
| *32* | Control | *Coreopsis tinctoria* | 4 | 1 | 1 | 1 | 66.67 | NA |
| *32* | Control | *Rudbeckia hirta* | 10 | 0 | 2 | 12 | 501 | NA |
| *33* | Clothianidin | *Coreopsis tinctoria* | 5 | 1 | 1 | 6 | 109 | 18.72 |
| *33* | Clothianidin | *Monarda fistulosa* | 1 | 0 | 0 | 0 | NA | 4.23 |
| *33* | Clothianidin | *Rudbeckia hirta* | 4 | 0 | 2 | 4 | 387.67 | 13.77 |
| *34* | Control | *Coreopsis tinctoria* | 7 | 2 | 2 | 5 | 117.67 | NA |
| *34* | Control | *Rudbeckia hirta* | 0 | 0 | 1 | 2 | 506.67 | NA |
| *35* | Clothianidin | *Coreopsis tinctoria* | 3 | 1 | 2 | 2 | 141.33 | 13.47 |

| | | | | | | | | |
|---|---|---|---|---|---|---|---|---|
| *35* | Clothianidin | *Monarda fistulosa* | 2 | 0 | 1 | 0 | NA | 21.43 |
| *35* | Clothianidin | *Rudbeckia hirta* | 1 | 0 | 0 | 3 | 751.67 | 11.64 |
| *37* | Clothianidin | *Coreopsis tinctoria* | 19 | 0 | 2 | 2 | 126 | 23.53 |
| *37* | Clothianidin | *Monarda fistulosa* | 1 | 0 | 0 | 1 | NA | 5.72 |
| *37* | Clothianidin | *Rudbeckia hirta* | 7 | 4 | 2 | 2 | 526 | 16.33 |

**Table S2:** Estimates, goodness of fit, and *p*-values for the homogeneity of variance analysis (*betadisper*) associated with our PERMANOVA.

| | ***df*** | **Sum of squares** | ***f*-value** | ***p*-value** |
|---|---|---|---|---|
| *Clothianidin* | 1 | 0.0007 | 0.086 | 0.771 |
| *Plant species* | 4 | 0.468 | 11.965 | **< 0.001** |

**Table S3:** Levels of clothianidin present in the flowers of collected plants from experimental plots. Clothianidin concentrations were standardized based on 100mg of plant tissue and parts per billion (ppb) concentrations were calculated.

| ***Treatment*** | ***Latin Name*** | ***Plot Number*** | ***Plant Part*** | ***ng per 100mg tissue*** | ***ppb*** |
|---|---|---|---|---|---|
| Clothianidin | *Chamaecrista fasciculata* | 9 | Flower | 8.4 | 83.99 |
| Clothianidin | *Chamaecrista fasciculata* | 13 | Flower | 0.3 | 3.12 |
| Clothianidin | *Chamaecrista fasciculata* | 17 | Flower | 1.9 | 18.90 |
| Clothianidin | *Coreopsis tinctoria* | 1 | Flower | 1.8 | 17.61 |
| Clothianidin | *Coreopsis tinctoria* | 9 | Flower | 1.6 | 15.54 |
| Clothianidin | *Coreopsis tinctoria* | 15 | Flower | 2.0 | 19.60 |
| Clothianidin | *Rudbeckia hirta* | 1 | Flower | 0.3 | 3.49 |
| Clothianidin | *Rudbeckia hirta* | 9 | Flower | 0.7 | 6.84 |
| Clothianidin | *Rudbeckia hirta* | 15 | Flower | 0.8 | 8.11 |
| Control | *Chamaecrista fasciculata* | 4 | Flower | < 0 | 0.00 |
| Control | *Chamaecrista fasciculata* | 6 | Flower | 0.2 | 1.79 |
| Control | *Chamaecrista fasciculata* | 12 | Flower | 0.6 | 6.06 |
| Control | *Coreopsis tinctoria* | 4 | Flower | 0.7 | 7.47 |
| Control | *Coreopsis tinctoria* | 16 | Flower | < 0 | 0.00 |
| Control | *Coreopsis tinctoria* | 20 | Flower | 1.1 | 10.98 |
| Control | *Rudbeckia hirta* | 2 | Flower | 0.4 | 4.08 |
| Control | *Rudbeckia hirta* | 4 | Flower | 0.5 | 5.34 |
| Control | *Rudbeckia hirta* | 10 | Flower | 4.5 | 45.33 |

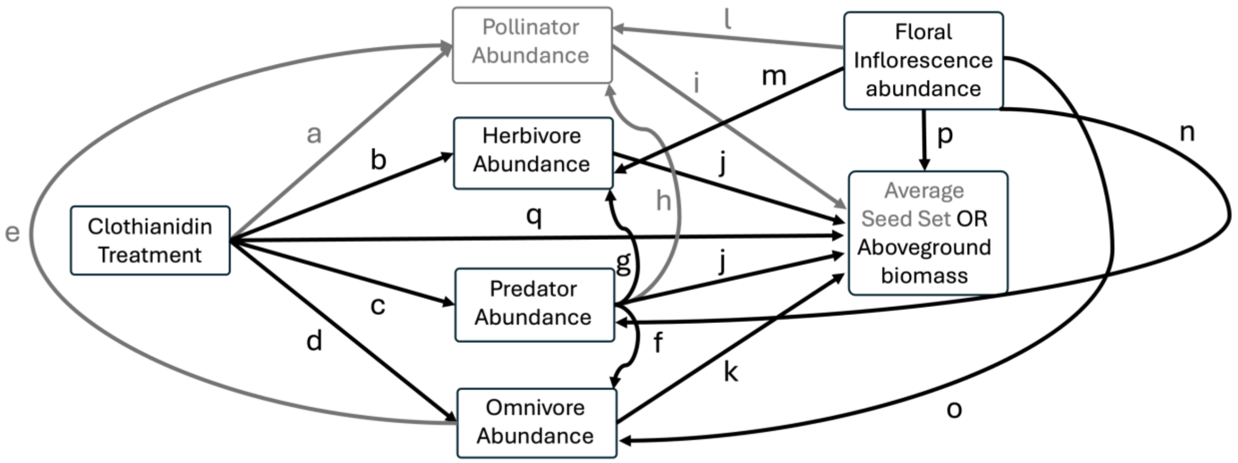


**Figure S1.** Conceptual representation and justification of the causal links in our hypothesized structural model. Boxes, letters, and paths in black were hypothesized for both seed set and aboveground biomass while gray connections represent only the seed set model. We expected that: (a) clothianidin treatment increases pollinator abundance (Kessler *et al.*, 2015; Arce *et al.*, 2018; Tetlie & Harmon-threatt, 2024); (b) clothianidin treatment decreases herbivore abundance (Harmon *et al.*, 2023; Lee & Davis, 2023); (c) clothianidin treatment decrease predator abundance (Tooming *et al.*, 2017; Harmon *et al.*, 2023); (d) clothianidin treatment decreases omnivore abundance (Uhl *et al.*, 2015; Schläppi *et al.*, 2020); (e) omnivore abundance decreases pollinator abundance (Junker *et al.*, 2007; Cembrowski *et al.*, 2014), (f) predator abundance decreases omnivore abundance (Hunter, 2009); (g) predator abundance decreases herbivore abundance (Moran & Hurd, 1997; Hunter, 2009); (h) predator abundance decreases pollinator abundance (Dukas, 2005; Romero *et al.*, 2011); (i) pollinator abundance increases average seed set (Garibaldi *et al.*, 2014; Cohen *et al.*, 2021); (j) herbivore abundance decreases seed set (Hendrix, 1988; Poveda *et al.*, 2003) and aboveground biomass (Nötzold *et al.*, 1997); (k) omnivore abundance increases seed set (Brown & Human, 1997); (l) floral inflorescence abundance increases pollinator abundance (Fowler *et al.*, 2016; Graystock *et al.*, 2020); (m) floral inflorescence abundance increases herbivore abundance (Rusman *et al.*, 2019); (n) floral inflorescence abundance increases predator abundance (Losapio *et al.*, 2016); (o) floral inflorescence abundance increases omnivore abundance (Matter *et al.*, 1999); floral inflorescence abundance decrases average seed set (Van Reeth *et al.*, 2019); and clothianidin treatment would increase average seed set (Wilde *et al.*, 2004; Zhang *et al.*, 2017).

### Literature Cited

Arce, A.N., Rodrigues, A.R., Yu, J., Colgan, T.J., Wurm, Y. & Gill, R.J. (2018) Foraging bumblebees acquire a preference for neonicotinoid-treated food with prolonged exposure. *Proceedings of the Royal Society B: Biological Sciences*, **285**, 8–11.

Brown, M.J.F. & Human, K.G. (1997) Effects of harvester ants on plant species distribution and abundance in a serpentine grassland. *Oecologia*, **112**, 237–243.

Cembrowski, A.R., Tan, M.G., Thomson, J.D. & Frederickson, M.E. (2014) Ants and ant scent reduce bumblebee pollination of artificial flowers. *American Naturalist*, **183**, 133–139.

Cohen, H., Philpott, S.M., Liere, H., Lin, B.B. & Jha, S. (2021) The relationship between pollinator community and pollination services is mediated by floral abundance in urban landscapes. *Urban Ecosystems*, **24**, 275–290.

Dukas, R. (2005) Bumble bee predators reduce pollinator density and plant fitness. *Ecology*, **86**, 1401–1406.

Fowler, R.E., Rotheray, E.L. & Goulson, D. (2016) Floral abundance and resource quality influence pollinator choice. *Insect Conservation and Diversity*, **9**, 481–494.

Garibaldi, L.A., Steffan-dewenter, I., Winfree, R., Aizen, M.A., Bommarco, R., Cunningham, S.A., *et al.* (2014) Wild Pollinators Enhance Fruit Set of Crops Regardless of Honey Bee Abundance. *Science*, **339**, 1608–1611.

Graystock, P., Ng, W.H., Parks, K., Tripodi, A.D., Muñiz, P.A., Fersch, A.A., *et al.* (2020) Dominant bee species and floral abundance drive parasite temporal dynamics in plant-pollinator communities. *Nature Ecology and Evolution*, **4**, 1358–1367.

Harmon, G.T., Harmon-Threatt, A.N. & Anderson, N.L. (2023) Changes in predator biomass may mask the negative effects of neonicotinoids on primary consumers in field settings. *Insect Conservation and Diversity*, **16**, 298–305.

Hendrix, S.D. (1988) Herbivory and its impact on plant reproduction. *Plant reproductive ecology: patterns and strategies*, 246–263.

Hunter, M.D. (2009) Trophic promiscuity, intraguild predation and the problem of omnivores. *Agricultural and Forest Entomology*, **11**, 125–131.

Junker, R., Chung, A.Y.C. & Blüthgen, N. (2007) Interaction between flowers, ants and pollinators: Additional evidence for floral repellence against ants. *Ecological Research*, **22**, 665–670.

Kessler, S.C., Tiedeken, E.J., Simcock, K.L., Derveau, S., Mitchell, J., Softley, S., *et al.* (2015) Bees prefer foods containing neonicotinoid pesticides. *Nature*, **521**, 74–76.

Lee, S.T. & Davis, J.A. (2023) Field and Forage Crops The impact of thiamethoxam on the feeding and behavior of 2 soybean herbivore feeding guilds, 1–15.

Losapio, G., Gobbi, M., Marano, G., Avesani, D., Boracchi, P., Compostella, C., *et al.* (2016) Feedback effects between plant and flower-visiting insect communities along a primary succession gradient. *Arthropod-Plant Interactions*, **10**, 485–495.

Matter, S.F., Landry, J.B., Greco, A.M. & Lacourse, C.D. (1999) Importance of floral phenology and florivory for Tetraopes tetraophthalmus (Coleoptera: Cerambycidae): Tests at the population and individual level. *Environmental Entomology*, **28**, 1044–1051.

Moran, M.D. & Hurd, L.E. (1997) A trophic cascade in a diverse arthropod community caused by a generalist arthropod predator. *Oecologia*, **113**, 126–132.

Nötzold, R., Blossey, B. & Newton, E. (1997) The influence of below ground herbivory and plant competition on growth and biomass allocation of purple loosestrife. *Oecologia*, **113**, 82–93.

Poveda, K., Steffan-Dewenter, I., Scheu, S. & Tscharntke, T. (2003) Effects of below- and aboveground herbivores on plant growth, flower visitation and seed set. *Oecologia*, **135**, 601–605.

Reeth, C. Van, Michel, N., Bockstaller, C. & Caro, G. (2019) Influences of oilseed rape area and aggregation on pollinator abundance and reproductive success of a co-flowering wild plant. *Agriculture, Ecosystems and Environment*, **280**, 35–42.

Romero, G.Q., Antiqueira, P.A.P. & Koricheva, J. (2011) A meta-analysis of predation risk effects on pollinator behaviour. *PLoS ONE*, **6**.

Rusman, Q., Karssemeijer, P.N., Lucas-Barbosa, D. & Poelman, E.H. (2019) Settling on leaves or flowers: herbivore feeding site determines the outcome of indirect interactions between herbivores and pollinators. *Oecologia*, **191**, 887–896.

Schläppi, D., Kettler, N., Straub, L., Glauser, G. & Neumann, P. (2020) Long-term effects of neonicotinoid insecticides on ants. *Communications Biology*, **3**, 1–9.
Tetlie, J. & Harmon-threatt, A. (2024) Neonicotinoid contamination in conservation areas affects bees more sharply than beetles. *Frontiers in Ecology and Evolution*, 1–11.
Tooming, E., Merivee, E., Must, A., Merivee, M.I., Sibul, I., Nurme, K., *et al.* (2017) Behavioural effects of the neonicotinoid insecticide thiamethoxam on the predatory insect Platynus assimilis. *Ecotoxicology*, **26**, 902–913.
Uhl, P., Bucher, R., Schäfer, R.B. & Entling, M.H. (2015) Sublethal effects of imidacloprid on interactions in a tritrophic system of non-target species. *Chemosphere*, **132**, 152–158.
Wilde, G., Roozeboom, K., Claassen, M., Janssen, K. & Witt, M. (2004) Seed treatment for control of early-season pests of corn and its effect on yield. *Journal of Agricultural and Urban Entomology*, **21**, 75–85.
Zhang, Z., Wang, Y., Zhao, Y., Li, B., Lin, J., Zhang, X., *et al.* (2017) Nitenpyram seed treatment effectively controls against the mirid bug Apolygus lucorum in cotton seedlings. *Scientific Reports*, **7**, 1–9.
